# Flatworms in the land of the midnight sun: first record of the invasive species *Obama nungara* (Platyhelminthes, Geoplanidae) in Sweden, its northernmost location in continental Europe

**DOI:** 10.1101/2025.03.24.644988

**Authors:** Chahinez Bouguerche, Jonas Roth, Jean-Lou Justine

## Abstract

The land flatworm *Obama nungara*, native to South America and already invasive in several European countries, is reported here for the first time in Sweden, marking its first sighting in Scandinavia. The finding of a specimen in Malmö in November 2024 suggests that *O. nungara* may be present in other areas of Sweden, with its establishment potentially dating back a certain time. The *cox*1 haplotype of the Malmö specimen clustered within a large group of haplotypes found in several countries of Europe; this group was found in previous studies to originate from Argentina. Within this group, the Malmö specimen belongs to the most common haplotype in Europe, which precludes any more precise assessment of its origin.

## Introduction

Invasive species are considered one of the major threats to biodiversity and records of invasive species in new territories are becoming increasingly numerous. Among invasive alien species, land flatworms (Platyhelminthes, Tricladida, Geoplanidae) are typical in that they are incapable of flying or swimming across oceans and that their dispersion is exclusively the consequence of human activities. In this particular case, the transport of potted plants is considered the major factor of land flatworm invasion in new territories (Sluys, 2016). Like other species, land flatworms can only thrive in conditions that are favourable to them, and climate is one of the major factors limiting their spread. Since land flatworms are generally associated with warm and humid climates (Sluys, 2016), a new record in a Northern European country is worth mentioning.

The flatworm *Obama nungara* Carbayo, Álvarez-Presas, Jones & Riutort, 2016 is a medium-sized land planarian with a lance-shaped body that can grow up to 70 millimetres in length. Its dorsal surface ranges in colour from golden to honey yellow and is patterned with dark brown to black spots and irregular flecks that form a marbled appearance, varying from light to dark brown. The ventral surface is smooth and uniformly coloured, typically cream to pale grey (Carbayo et al., 2016). *Obama nungara* is the most abundant land planarian in France (Justine et al., 2020) and is also present in a number of other European countries. It is a predator of earthworms and molluscs and is therefore a potential threat to the biodiversity of soils. It is able to proliferate at high rates in gardens in temperate climates, with thousands of specimens in a single garden (Justine et al., 2020; Noël et al., 2025). Its precise dietary preferences are known with precision for earthworms in France (Roy et al., 2022), but our level of knowledge on its mollusc diet is still rudimentary. The predation of earthworms by this predatory flatworm, and the subsequent disruption of native soil communities, can significantly impact soil functionality. Earthworms, often referred to as ecosystem engineers, play a crucial role in maintaining soil functions and delivering essential ecosystem services.

We report here the first mention of *Obama nungara* in Sweden, marking its inaugural record in Scandinavia and setting off an important alarm for the region. This finding underscores the significance of monitoring invasive flatworms in Sweden, as their spread can have considerable ecological implications. Early detection and ongoing surveillance are crucial to preventing further establishment and potential negative impacts on local ecosystems.

## Material and Methods

### Collection of specimens

One of us (JR) was in Malmö, Sweden conducting a search for foreign snails and slugs in a public garden (Monday 4 November 2024, in the evening around 22.00; Coordinates: 55°35’55.7”N 12°59’55.1”E) on behalf of the County Administrative Board (Roth, 2025). He observed a yellowish animal moving along the ground and immediately photographed and collected the live specimen. Further inspection of the area revealed four additional flatworms, discovered within approximately 15 minutes. Specimens were fixed in ethanol and registered into the Invertebrates collection of the Swedish Museum of Natural History (SMNH), Stockholm, Sweden as XXX.

### Molecular methods, sequencing and haplotype network

A small part was excised from one specimen and subjected to Sanger sequencing of the partial *cox1* gene, following a routine protocol as defined by Justine et al. (2020). Genomic DNA was extracted using a QIAamp DNA Mini Kit (QIAGEN, Hilden, Germany). Two sets of primers were used to amplify the *cox*1 gene. A fragment of 424 bp (designated in this text as ‘short sequence’) was amplified with the primers JB3 (=COI-ASmit1) (forward 5′-TTTTTTGGGCATCCTGA GGTTTAT-3′) and JB4.5 (=COI-ASmit2) (reverse 5′-TAAAGAAAGAACATAATGA AAATG-3′) (Bowles et al., 1995; Littlewood et al., 1997). PCR reactions were performed in 20 mL, containing 1 ng of DNA, 1Å∼ CoralLoad PCR buffer, three mM MgCl2, 66 μm of each dNTP, 0.15 mm of each primer and 0.5 units of Taq DNA polymerase (QIAGEN). The amplification protocol was: 4′ at 94°C, followed by 40 cycles of 94°C for 30″, 48°C for 40″, and 72°C for 50″, with a final extension at 72°C for 7′. A fragment of 825 bp was amplified with the primers BarS (forward 5′-GTTATGCCTGTAATGATTG-3′) (Álvarez-Presas et al., 2011) and COIR (reverse 5′-CCWGTYARMCCHCCWAYAGTAAA-3′) (Lázaro et al., 2009; Mateos et al., 2013). PCR products were purified and sequenced in both directions on a 96-capillary 3730xl DNA analyser sequencer (Applied Biosystems, Foster City, CA, USA). Results of both analyses were concatenated to obtain a COI sequence of 909 bp in length (designated in this text as ‘long sequence’). Sequences were edited using CodonCode Aligner software (CodonCode Corporation, Dedham, MA, USA), compared to the GenBank database content using BLAST, and deposited in GenBank under Accession Number xxxx. Nomenclature of *O. nungara* clades follows Justine et al. (2020).

### Matrix

We used a matrix made public by Justine et al. (2020) to which we added a sequence of *O. nungara* from La Réunion (Justine et al., 2022) and the new sequence obtained from the Swedish specimen. The matrix is 255 bases in length and includes 99 sequences, namely sequences of seven *Obama* species: *O. anthropophila, O. burmeisteri, O. carinata, O. josefi, O. ladislavii, O. maculipunctata*, and *O. marmorata*) and 92 sequences of *O. nungara*, including our new sequence. The matrix includes sequences of *O. nungara* from Argentina, Brazil, and eight territories in Europe.

### Trees and distances

The alignment was constructed separately in AliView (Larsson, 2014) and trimmed to the shortest sequence. Nucleotide substitution models for phylogenetic analyses using the maximum likelihood (ML) method were estimated using MEGA11 (Stecher et al., 2020). The Hasegawa-Kishino-Yano model (Hasegawa et al., 1985) with invariant sites (HKY+I) was used, with 500 bootstraps. The neighbour-joining (NJ) method (Saitou & Nei, 1987) was also used for comparison in MEGA11, with 2 000 bootstraps. p-distances and the Kimura two-parameter (K2P) distances (Kimura, 1980) were computed from the same datasets with MEGA11. Trees were constructed in MEGA11.

### Population analysis

The haplotype network was constructed using a matrix made public (Justine et al., 2020) and adding a sequence from La Réunion (Justine et al., 2022) and the new sequence obtained from the Swedish specimen. In the nexus file, traits were the geographical origin of the species, with number of traits set to 10, and trait labels set as: Argentina Brazil, UK (including Guernsey), Portugal, Spain, Italy, Switzerland, Belgium, Metropolitan France, La Réunion, and Sweden. The matrix had 93 taxa and was 255 bases in length. We used PopART (Leigh & Bryant, 2015) to create the haplotype network.

## Results

### Morphology

Order Tricladida Lang, 1884.

Family Geoplanidae Stimpson, 1857

Subfamily Geoplaninae Stimpson, 1857

*Obama* Carbayo, Álvarez-Presas, Olivares, Marques, Froehlich & Riutort, 2013

*Obama nungara* Carbayo, Álvarez-Presas, Jones & Riutort, 2016

### Short description (Figure 1)

Swedish specimens, preserved in ethanol: leaf-like body, total length 3.5–4.8 cm, 0.5–0.7 cm wide. Eyes on a single row at the front margin and multiple rows along the sides further back. Dorsal surface with marbled pattern in varying shades of brown, nearly black **(Fig 1 A-D)**. Relative mouth/body length 61%, relative gonopore/body length 75% (measured on 1 flat specimen). Colour pattern of live specimens, orange dorsal colour **(Fig 1 E)** and beige ventral colour **(Fig 1 F)**. Among the specimens observed live, one was observed feeding on an unidentified earthworm, which displayed abnormal movement.

**Figure 1.**
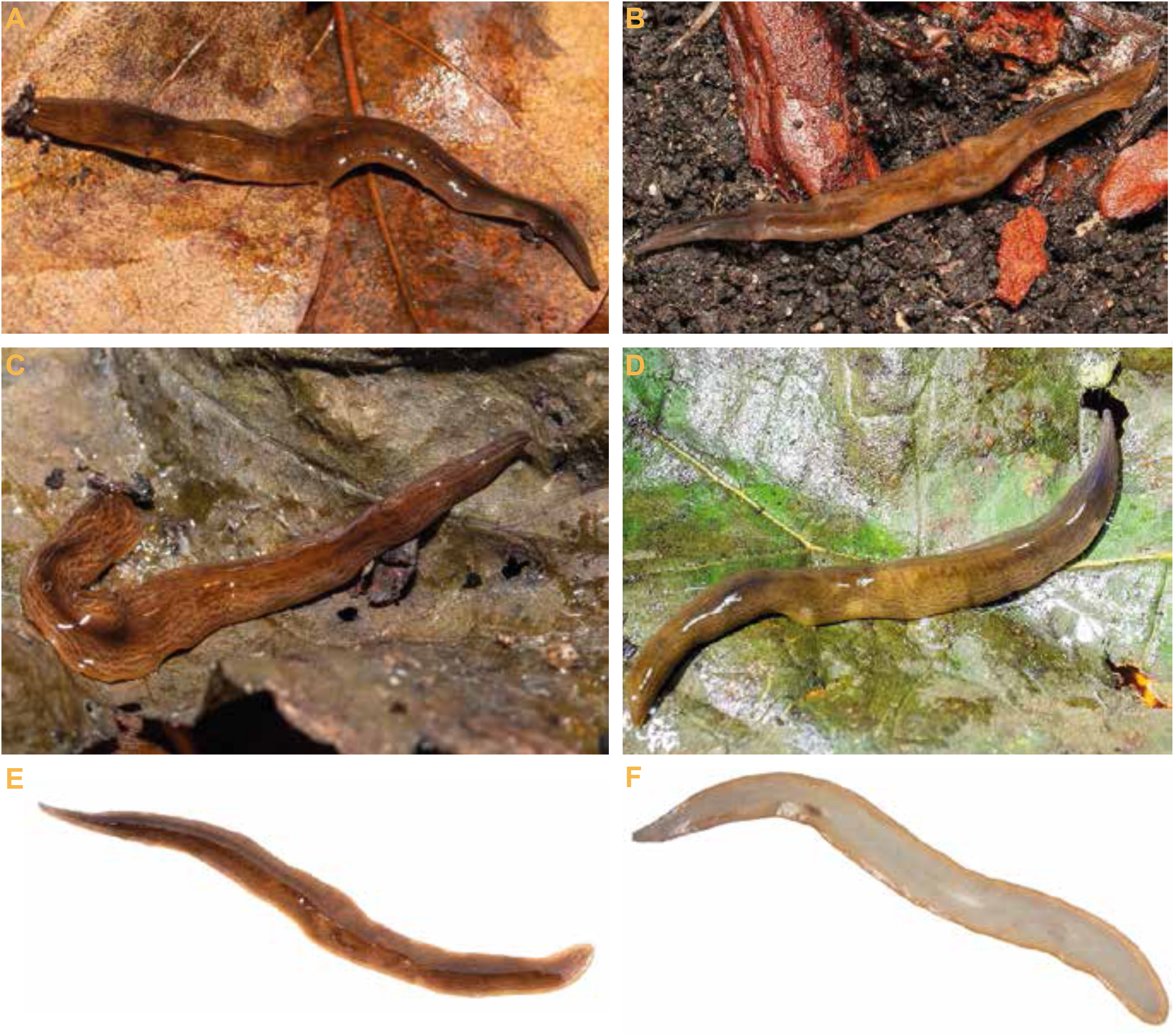
*Obama nungara*, Specimen from Malmö, Sweden. Photo by Jonas Roth. A, dorsal view on a leaf, exhibiting colour mimicry. B, dorsal view of the worm in its natural habitat. This specimen shows dorsal ‘tiger stripes’. C, on a leaf, anterior part showing eyes. D, nocturnal activity of the worm, observed under flashlight, examining plants and growing areas. E, specimen showing dorsal side. F, specimen showing ventral side. Note that the ventral surface is remarkably pale compared to the coloured dorsal side.

The colour pattern, overall body shape, and body length ratios are all consistent with the specific identification of the Malmö specimen as *O. nungara*.

### Molecular analysis

The partial *cox*1 sequence of the Malmö specimen (GenBank number XXX) was identical (100% identity) to several sequences from various European countries and a sequence from Argentina.

A phylogenetic tree **(Fig. 2)** was constructed using a matrix of *Obama nungara cox*1 sequences from both GenBank and newly generated data, with sequences from other species serving as the outgroup. The *O. nungara* clade, consisted of 92 sequences, with strong nodal support (ML 91%, NJ 99%). Due to the species’ complex nomenclatural history, several GenBank entries labelled as *O. marmorata* or *Obama* sp. were included within this clade, and we consider them representatives of *O. nungara*.

**Figure 2.**
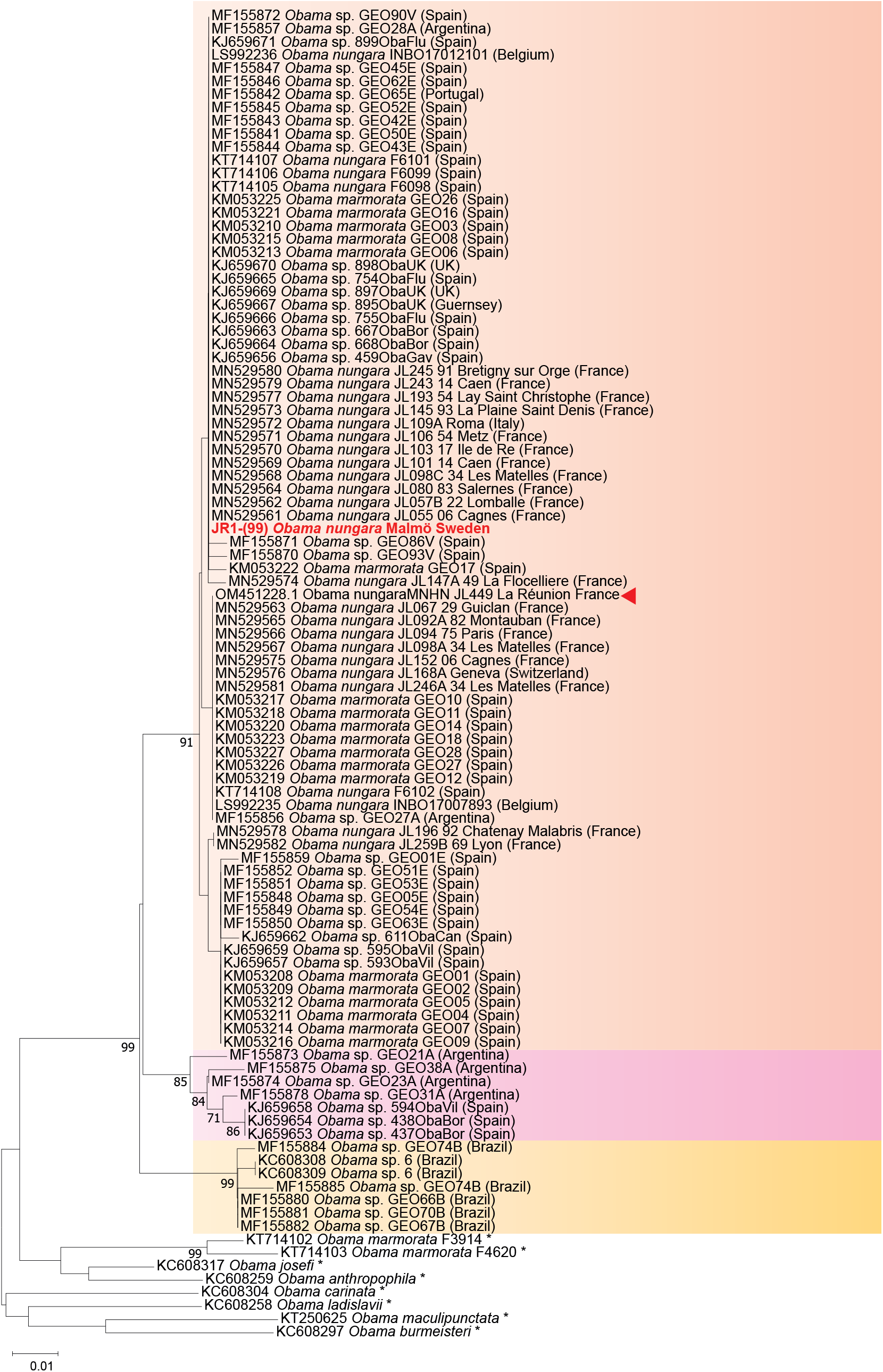
Maximum-likelihood tree of *Obama nungara* and its close relatives. The analysis included 101 nucleotide sequences with a total of 255 positions in the final dataset. The matrix incorporated selected sequences of *O. nungara* along with six other *Obama* species (marked with *) following Justine et al. (2020). The percentage of trees supporting each clade is indicated next to the branches (ML: maximum likelihood; NJ: neighbour-joining). The geographic origin of each sequence is specified. The newly generated sequence from Sweden is highlighted in red, while the red arrow marks the *O. nungara* sequence from La Réunion, France. Notably, the *O. nungara* clade contains multiple sequences in GenBank under different names.

A simplified version of the phylogenetic tree, focusing solely on the *O. nungara* clade, is presented in **Figure 3**. Within this clade, three well-supported subclades were identified. The ‘Argentina 1’ clade (ML 87%, NJ 91%) is the most robust and encompasses the majority of *O. nungara* sequences, including all specimens from Sweden and other European countries, along with specimens from Argentina. This clade forms a sister group to the ‘Argentina 2’ clade (ML 80%, NJ 85%), which consists exclusively of specimens from Argentina and Spain. The ‘Brazil’ clade (ML 88%, NJ 99%) includes only specimens from Brazil.

**Figure 3.**
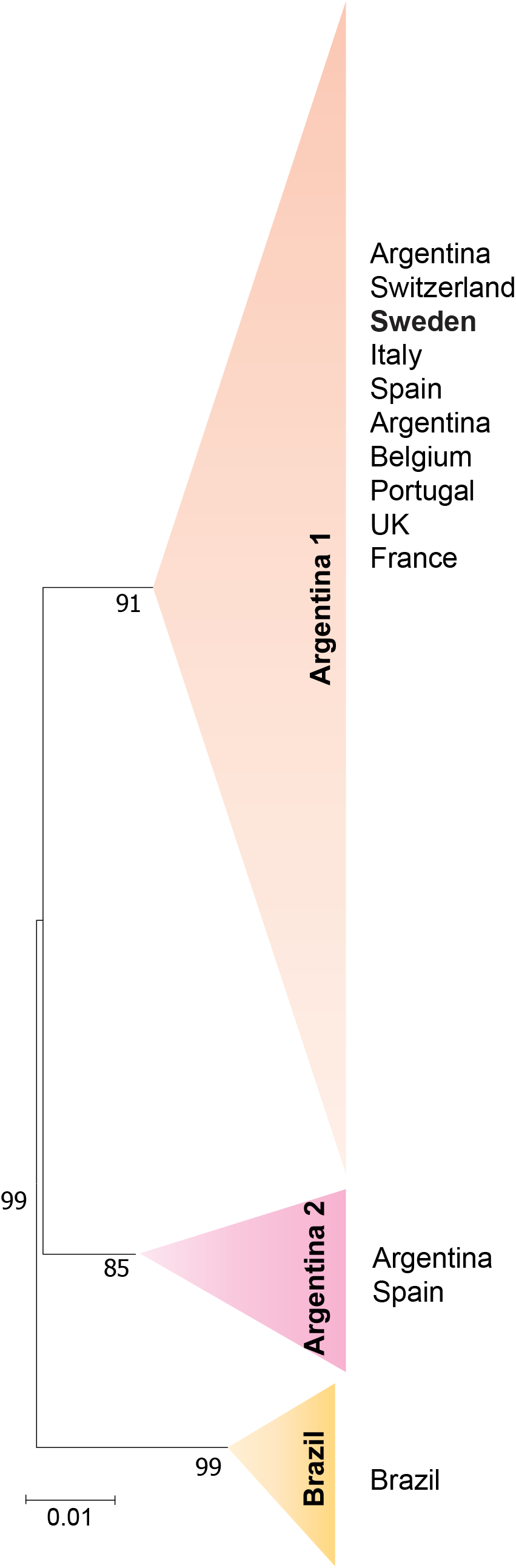
Simplified phylogenetic tree showing the relationships within *Obama nungara* and its closely related species showing only the *Obama nungara* clade. This tree is derived from the complete tree presented in **Figure 2**. *Obama nungara* is divided into three well-supported clades, representing distinct populations: ‘Argentina 1,’ ‘Argentina 2,’ and ‘Brazil.’ Invasive specimens from several European countries—Spain, Portugal, France, the UK, Italy, and Switzerland—are placed within the ‘Argentina 1’ clade. The haplotype of *Obama nungara* from Sweden clusters within the ‘Argentina 1’ group, which also includes sequences from Argentina and other invaded European countries.

The haplotype network is presented in **Figure 4**, revealing 19 haplotypes for *O. nungara*. The analysis identified three main networks. Network A, the largest in terms of sequence numbers, comprised 10 haplotypes and included specimens from Argentina and eight European countries: Spain, Portugal, France, Belgium, the UK, Italy, Switzerland, and Sweden. This network corresponded to the ‘Argentina 1’ clade in the phylogenetic tree. Within Network A, three major haplotypes exhibited distinct geographic distributions. The largest haplotype contained representatives from Argentina and multiple European countries, including the Malmö specimen. The second haplotype included specimens from Argentina and Europe, while the third and smaller one was exclusive to Spain. Network B consisted of five haplotypes and corresponded to the ‘Argentina 2’ clade in the tree. It included four haplotypes from Argentina and one from Spain. Network C, comprising four haplotypes, included only specimens from Brazil and aligned with the ‘Brazil’ clade in the phylogenetic tree.

**Figure 4.**
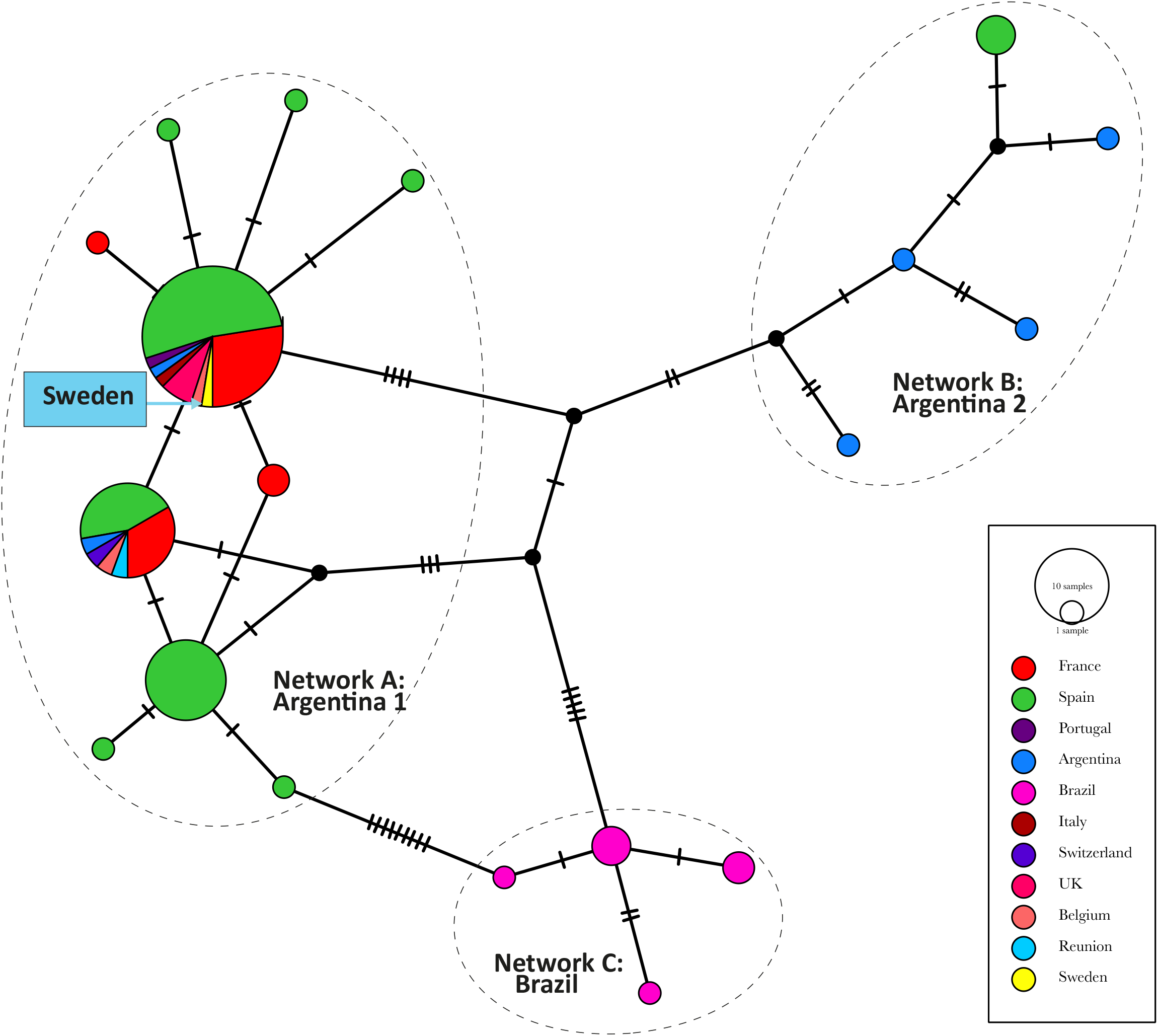
Network of relationships between specimens of *Obama nungara* from various localities. Three distinct groups are observed: ‘Argentina 1’, ‘Argentina 2’, and ‘Brazil’. The haplotype of *Obama nungara* from Sweden clusters within the primary group, ‘Argentina 1’ which comprises sequences from Argentina as well as all European invaded countries. The number of mutations (nucleotide differences) between haplotypes is indicated via black lines with putative haplotypes depicted as black-filled circles.

## Discussion

As mentioned above, most invasive land planarians come from warm climates; representative examples are *Bipalium kewense* Moseley, 1878, now almost cosmopolitan in near-tropical areas of the world and even the warmer parts of Europe, including Southern France, Italy, Spain and Portugal (Justine et al., 2018; Winsor, 1983), or *Platydemus manokwari* de Beauchamp, 1963, found in many islands and territories in warm areas of the Pacific and recently in the southern United States, especially Florida (Justine et al., 2015). However, some species are more adapted to colder climate. A notable example is *Arthurdendyus triangulatus* (Dendy, 1896) Jones, 1999, originating from the southern regions of New Zealand: the species has invaded the British Isles, but is restricted to their most northern parts (Murchie & Gordon, 2013). This species is currently the only geoplanid included in the list of Invasive Alien Species of Union concern (European_and_Mediterranean_Plant_Protection_Organization, 2000). Another example is *Bipalium adventitium* Hyman, 1943, for which the origin is unknown, but which can survive the snowy winters of northern USA and even Canada (Justine et al., 2019); a modelling study predicted that it could invade most of continental Europe including part of the Scandinavian peninsula (Fourcade et al., 2022).

*Obama nungara* originates from a region at the boundary of Southern Brazil and Argentina (Carbayo et al., 2016), in which the climate is not warmer than in many parts of Europe. The species has now been found in most European countries (de Waart et al., 2025; Justine et al., 2020). In 2020, Justine *et al*. wrote that the species had never been recorded “in Germany or any country east of Germany” (Justine et al., 2020). Since then, this statement has become inaccurate and several records were added for Germany (Glaw et al., 2024), Hungary (Lazanyi et al., 2024), and Slovakia (Čapka & Čejka, 2021). Records have also recently been reported for the Netherlands (de Waart et al., 2025). In the Mediterranean region, the species has invaded islands such as Corsica (Justine et al., 2020) and Malta (Cilia, 2024). A record from La Réunion, an island in the Indian Ocean off the African coast, was attributed to transport from Metropolitan France, possibly with stones used for construction (Justine et al., 2022); all records in Europe are thought to result from transport of potted plants from country to country. The presence of *O. nungara* in the USA is possible in view of citizen science records, but has not yet been confirmed by molecular identification (Justine et al., 2024).

A modelling study, based on available records of the species both in its native habitat and invaded areas, proposed suitability maps for *O. nungara* (Fourcade, 2021). Under the current climate, the southern part of Sweden is included in areas where *O. nungara* could live, however with low suitability. Therefore, the finding in Malmö reported in the present paper is not aberrant, and more records from the Scandinavian peninsula, at least the southern parts of Sweden and Norway, are to be expected in the future. The specimens in Malmö were found in autumn; it remains to be seen whether records will continue after winter.

*Obama nungara* is a generalist predator, preying on earthworms, terrestrial molluscs (snails, slugs) and other land planarians (Boll et al., 2015; Carbayo et al., 2016; Justine et al., 2020; Roy et al., 2022). Its invasion in Sweden could thus be a threat to soil biodiversity.

The 100% similarity of the *cox1* sequence of the Swedish specimen with sequences from several countries of Europe precludes any precise assessment of its geographical origin based on our molecular data, which are limited to a portion of a single gene. The Swedish media reported that the public garden in Malmö was filled with plants imported from Germany (Sweden_Herald, 2025), and this is an acceptable possibility as much as any other.

Recent observations indicate that *O. nungara* has successfully overwintered and survived sub-zero temperatures by seeking microhabitats with relatively stable and warmer conditions, such as beneath rocks and decomposing plant material. The County Administrative Board has expressed concern regarding the potential spread of *O. nungara* in Sweden during the spring. This risk is heightened by the increased trade and transportation of plants from nurseries, as soil and plant material may inadvertently facilitate the dispersal of the species into new environments (Rikare_Trädgård, 2025).

An unexpected positive outcome of this finding is that, after media in Sweden made the finding of *Obama nungara* known by the public (Anonymous, 2024, 2024b; Nystrand, 2024; Stad, 2024), we received several reports of other species of land flatworm from Sweden, sent by scientists or concerned citizens. Most of these records, however, are from hothouses, and, in spite of their scientific interest, do not have the same biogeographical and ecological interest as the finding in the open in Malmö.

## Conclusion

The discovery of *Obama nungara* in Malmö aligns with modelling predictions that parts of southern Sweden could provide a suitable habitat for the species, albeit with low suitability. Given its adaptability and previous invasion history, further records from Sweden and neighbouring Scandinavian countries are likely. The species’ potential impact on local soil biodiversity, due to its generalist predatory nature, underscores the need for monitoring and further research. Available genetic data show that the Malmö specimen belongs to a haplotype widespread in Europe. A noteworthy outcome of this discovery is the increased public awareness of invasive land planarians in Sweden, leading to additional reports of land flatworms, although many were confined to hothouses. This highlights the importance of citizen science in tracking and understanding the spread of invasive species.

## References

Álvarez-Presas, M., Carbayo, F., Rozas, J., & Riutort, M. (2011). Land planarians (Platyhelminthes) as a model organism for fine-scale phylogeographic studies: understanding patterns of biodiversity in the Brazilian Atlantic Forest hotspot. Journal of Evolutionary Biology, 24(4), 887–896. 10.1111/j.1420-9101.2010.02220.x

Anonymous. (2024). Invasiva plattmasken Obama nungara upptäckt i Skåne. https://www.lansstyrelsense/skane/om-oss/nyheter-och-press/nyheter---skane/2024-11-12-invasiva-plattmaskenobama-nungara-upptackt-i-skanehtml

Anonymous. (2024b). Invasive cannibal flatworm found in Malmö. Sweden Herald:https://swedenherald.com/article/invasive-cannibal-flatworm-found-in-malmo

Boll, P. K., Rossi, I., Amaral, S. V., & Leal-Zanchet, A. (2015). A taste for exotic food: Neotropical land planarians feeding on an invasive flatworm. PeerJ, 3, e1307. 10.7717/peerj.1307

Bowles, J., Blair, D., & McManus, D. P. (1995). A molecular phylogeny of the human schistosomes. Molecular Phylogenetics and Evolution, 4(2), 103–109. 10.1006/mpev.1995.1011

Capka, J., & Čejka, T. (2021). First record of Obama nungara in Slovakia (Platyhelminthes: Geoplanidae). Biodiversity & Environment, 13(2), 41-44.

Carbayo, F., Alvarez-Presas, M., Jones, H. D., & Riutort, M. (2016). The true identity of Obama (Platyhelminthes: Geoplanidae) flatworm spreading across Europe. Zoological Journal of the Linnean Society, 177, 5–28. 10.1111/zoj.12358

Cilia, D. (2024). On the occurrence of the invasive terrestrial flatworm Obama nungara Carbayo, Álvarez-Presas, Jones & Riutort, 2016 (Platyhelminthes Geoplanidae) in the Maltese islands. Naturalista Siciliano 48:, 35–40. 10.5281/zenodo.12075407

de Waart, S., Vanhove, M., Kmentová, N., & J-L., J. (2025). Going Dutch: Invasion pathways and current European distribution of non-native land flatworm species belonging to Geoplaninae and Bipaliinae with focus on the Netherlands. ARPHA Preprints 10.3897/arphapreprints.e146092

European_and_Mediterranean_Plant_Protection_Organization. (2000). EPPO Standards. Guidelines on Arthurdendyus triangulatus. Import requirements concerning Arthurdendyus triangulatus. http://archives.eppo.int/EPPOStandards/PM1_GENERAL/pm1-03-e.doc.

Fourcade, Y. (2021). Fine-tuning niche models matters in invasion ecology. A lesson from the land planarian Obama nungara. Ecological Modelling 457, 109686. 10.1016/j.ecolmodel.2021.10968

Fourcade, Y., Winsor, L., & Justine, J.-L. (2022). Hammerhead worms everywhere? Modelling the invasion of bipaliin flatworms in a changing climate. Diversity and Distributions 28, 844–858. 10.1111/ddi.13489

Glaw, F., Mass, R., Glaw, T., & Schreiner, J. (2024). Non-native terrestrial planarian species in Germany and Austria, with first locality records of Caenoplana variegata for both countries. SPIXIANA, 47, 113–118.

Hasegawa, M., Iida, Y., Yano, T., Takaiwa, F., & Iwabuchi, M. (1985). Phylogenetic relationships among eucaryotic kingdoms inferred from ribosomal RNA sequences. Journal of Molecular Evolution, 22, 32–38.

Justine, J.-L., Gastineau, R., Gey, D., Robinson, D., Bertone, M., & Winsor, L. (2024). A new species of alien flatworm in the Southern United States. PeerJ 12, e17904. 10.7717/peerj.17904

Justine, J.-L., Marie, A., Gastineau, R., Fourcade, Y., & Winsor, L. (2022). The invasive land flatworm Obama nungara in L. Réunion, a French island in the Indian Ocean, the first report of the species for Africa. Zootaxa 5154 469–476. 10.11646/zootaxa.5154.4.4

Justine, J.-L., Thery, T., Gey, D., & Winsor, L. (2019). First record of the invasive land flatworm Bipalium adventitium (Platyhelminthes, Geoplanidae) in Canada. Zootaxa, 4656(3), 591–595.

Justine, J.-L., Winsor, L., Barrière, P., Fanai, C., Gey, D., Han, A. W. K., La Quay-Velazquez, G., Lee, B. P. Y.-H., Lefevre, J.-M., Meyer, J.-Y., Philippart, D., Robinson, D. G., Thévenot, J., & Tsatsia, F. (2015). The invasive land planarian Platydemus manokwari (Platyhelminthes, Geoplanidae): records from six new localities, including the first in the USA. PeerJ, 3, e1037. 10.7717/peerj.1037

Justine, J.-L., Winsor, L., Gey, D., Gros, P., & Thévenot, J. (2018). Giant worms chez moi! Hammerhead flatworms (Platyhelminthes, Geoplanidae, Bipalium spp., Diversibipalium spp.) in metropolitan France and overseas French territories. PeerJ, 6, e4672. 10.7717/peerj.4672

Justine, J.-L., Winsor, L., Gey, D., Gros, P., & Thévenot, J. (2020). Obama chez moi! The invasion of metropolitan France by the land planarian Obama nungara (Platyhelminthes, Geoplanidae). PeerJ, 8, e8385.

Kimura, M. (1980). A simple method for estimating evolutionary rates of base substitutions through comparative studies of nucleotide sequences. Journal of Molecular Evolution, 16(2), 111–120.

Larsson, A. (2014). AliView: a fast and lightweight alignment viewer and editor for large data sets. Bioinformatics 30, 3276–3278.

Lazanyi, E., Boll, P., Pall-Gergely, B., Simon, J., Szeder, K., Turoci, A., & Katona, G. (2024). First records of alien land planarians (Platyhelminthes: Geoplanidae) in Hungary. Zootaxa 5403, 592–596 10.11646/zootaxa.5403.5.6

Lázaro, E. M., Sluys, R., Pala, M., Stocchino, G. A., Baguñà, J., & Riutort, M. (2009). Molecular barcoding and phylogeography of sexual and asexual freshwater planarians of the genus Dugesia in the Western Mediterranean (Platyhelminthes, Tricladida, Dugesiidae). Molecular Phylogenetics and Evolution, 52(3), 835–845. 10.1016/j.ympev.2009.04.022

Leigh, J. W., & Bryant, D. (2015). POPART: full-feature software for haplotype network construction. Methods in Ecology and Evolution, 6(9), 1110–1116. 10.1111/2041-210x.12410

Littlewood, D. T. J., Rohde, K., & Clough, K. A. (1997). Parasite speciation within or between host species? - Phylogenetic evidence from site-specific polystome monogeneans. International Journal for Parasitology, 27, 1289–1297. 10.1016/S0020-7519(97)00086-6

Mateos, E., Tudó, A., Álvarez-Presas, M., & Riutort, M. (2013). Planàries terrestres exòtiques a la Garrotxa. Annals de la Delegació de la Garrotxa de la Institució Catalana d’Història Natural, 6, 67–73.

Murchie, A. K., & Gordon, A. W. (2013). The impact of the “New Zealand flatworm”, Arthurdendyus triangulatus, on earthworm populations in the field. Biological Invasions, 15(3), 569–586. 10.1007/s10530-012-0309-7

Noël, S., Fourcade, Y., Roy V., Bonnet, G., and Dupont, L. (2025). Population Dynamics of the Exotic Flatworm Obama nungara in an Invaded Garden. Ecology and Evolution, 15(1), e70827. 10.1002/ece3.70827

Nystrand, L. (2024). Köttätande maskar sprider sig i Malmös planteringar. SVT Nyheter:https://www.svt.se/nyheter/lokalt/skane/kottatande-maskar-sprider-sig-i-malmos-planteringar

Rikare_Trädgård. (2025). Lövplattmasken överlevde vintern – se upp inför trädgårdssäsongen. Rikare Trädgård. https://rikaretradgard.se/lovplattmasken-overlevde-vintern-se-upp-infor-tradgardssasongen/.

Roth, J. (2025). Inventering av landlevande mollusker i Malmö – med fokus på främmande arter. Available from: https://www.lansstyrelsen.se/skane/om-oss/varatjanster/publikationer/2025/inventering-av-landlevande-mollusker-i-malmo---medfokus-pa-frammande-arter.html

Roy, V., Ventura, M., Fourcade, Y., Justine, J.-L., Gigon, A., & Dupont, L. (2022). Gut content metabarcoding and citizen science reveal the earthworm prey of the exotic terrestrial flatworm, Obama nungara. European Journal of Soil Biology 113, 103449. 10.1016/j.ejsobi.2022.103449

Saitou, N., & Nei, M. (1987). The neighbor-joining method: a new method for reconstructing phylogenetic trees. Molecular Biology and Evolution, 4, 406–425.

Sluys, R. (2016). Invasion of the Flatworms. American Scientist, 104(5), 288–295.

Stad, M. (2024). Plattmask från Sydamerika upptäckt i plantering. https://malmose/Aktuellt/Artiklar-Malmo-stad/2024-11-13-Plattmask-fran-Sydamerika-upptackt-i-planteringhtml

Stecher, G., Tamura, K., & Kumar, S. (2020). Molecular Evolutionary Genetics Analysis (MEGA) for macOS. Molecular Biology and Evolution 37, 1237–1239. 10.1093/molbev/msz312

Sweden_Herald. (2025). Invasive cannibal flatworm found in Malmö. Sweden Herald. https://swedenherald.com/article/invasive-cannibal-flatworm-found-in-malmo.

Winsor, L. (1983). A revision of the Cosmopolitan land planarian Bipalium kewense Moseley, 1878 (Turbellaria: Tricladida: Terricola). Zoological Journal of the Linnean Society, 79, 61–100. 10.1111/j.1096-3642.1983.tb01161.x

